# Augmented Reality Powers a Cognitive Prosthesis for the Blind

**DOI:** 10.1101/321265

**Authors:** Yang Liu, Noelle R. B. Stiles, Markus Meister

## Abstract

To restore vision for the blind several prosthetic approaches have been explored that convey raw images to the brain. So far these schemes all suffer from a lack of bandwidth and the extensive training required to interpret unusual stimuli. Here we present an alternate approach that restores vision at the cognitive level, bypassing the need to convey sensory data. A wearable computer captures video and other data, extracts the important scene knowledge, and conveys that through auditory augmented reality. This system supports many aspects of visual cognition: from obstacle avoidance to formation and recall of spatial memories, to long-range navigation. Neither training nor modification of the physical environment are required: Blind subjects can navigate an unfamiliar multi-story building on their first attempt. The combination of unprecedented computing power in wearable devices with augmented reality technology promises a new era of non-invasive prostheses that are limited only by software.

**Impact Statement:** A non-invasive prosthesis for blind people endows objects in the environment with voices, allowing a user to explore the scene, localize objects, and navigate through a building with minimal training.

## Introduction

About 36 million people are blind worldwide (Bourne et al., 2017). In industrialized nations, the dominant causes of blindness are age-related diseases of the eye, all of which disrupt the normal flow of visual data from the eye to the brain. In some of these cases biological repair is a potential option, and various treatments are being explored involving gene therapy, stem cells, or transplantation (Scholl et al., 2016). However, the dominant strategy for restoring vision has been to bring the image into the brain’s visual system through alternate means. The most direct route is electrical stimulation of surviving cells in the retina (Stingl and Zrenner, 2013; Weiland and Humayun, 2014) or of neurons in the visual cortex (Dobelle et al., 1974). Another option involves translating the raw visual image into a different sensory modality (Loomis et al., 2012; Proulx et al., 2016), such as touch (Stronks et al., 2016) or hearing (Auvray et al., 2007; Capelle et al., 1998; Meijer, 1992). So far, none of these approaches has enabled any practical recovery of the functions formerly supported by vision. Despite decades of efforts all users of such devices remain legally blind (Luo and da Cruz, 2016; Stingl et al., 2017; Striem-Amit et al., 2012; Stronks et al., 2016). While one can certainly hope for progress in these domains, it is worth asking what are the fundamental obstacles to current visual prostheses.

The human eye takes in about 1 gigabit of raw image information every second, whereas our visual system extracts from this just tens of bits to guide our thoughts and actions (Pitkow and Meister, 2014). All the above prosthetic approaches seek to transmit the raw image into the brain. This requires inordinately high data rates. Further, the signal must arrive in the brain in a format that can be interpreted usefully by the visual system or some substitute brain area to perform the key steps of knowledge acquisition, like scene recognition and object identification. None of the technologies available today deliver the high data rate required to retain the relevant details of a scene, nor do they produce a neural code for the image information that matches the capabilities and expectations of the human brain.

Three decades ago, one of the pioneers of sensory substitution articulated his vision of a future visual prosthesis (Collins, 1985): “I strongly believe that we should take a more sophisticated approach, utilizing the power of artificial intelligence for processing large amounts of detailed visual information in order to substitute for the missing functions of the eye and much of the visual pre-processing performed by the brain. We should off-load the blind travelers’ brain of these otherwise slow and arduous tasks which are normally performed effortlessly by the sighted visual system”. Whereas at that time the goal was hopelessly out of reach, today’s capabilities in computer vision, artificial intelligence, and miniaturized computing power are converging to make it realistic. Here we present such an approach that bypasses the need to convey the sensory data entirely, and focuses instead on the important high-level knowledge, presented at a comfortable data rate and in an intuitive format.

## Results

### Design principles

The new system is based on the Microsoft HoloLens (Fig. 1A), a powerful head-mounted computer designed for augmented reality (Hoffman, 2016). The HoloLens scans all surfaces in the environment using video and infrared sensors, creates a 3D map of the space, and localizes itself within that volume to a precision of a few centimeters (Fig. S1). It includes a see-through display for digital imagery superposed on the real visual scene; open ear speakers that augment auditory reality while maintaining regular hearing; and an operating system that implements the localization functions and provides access to the various sensor streams. We designed applications using the Unity game development platform which allows tracking of the user’s head in the experimental space; the simulation of virtual objects; the generation of speech and sounds that appear to emanate from specific locations; and interaction with the user via voice commands and a clicker.

**Figure 1.**
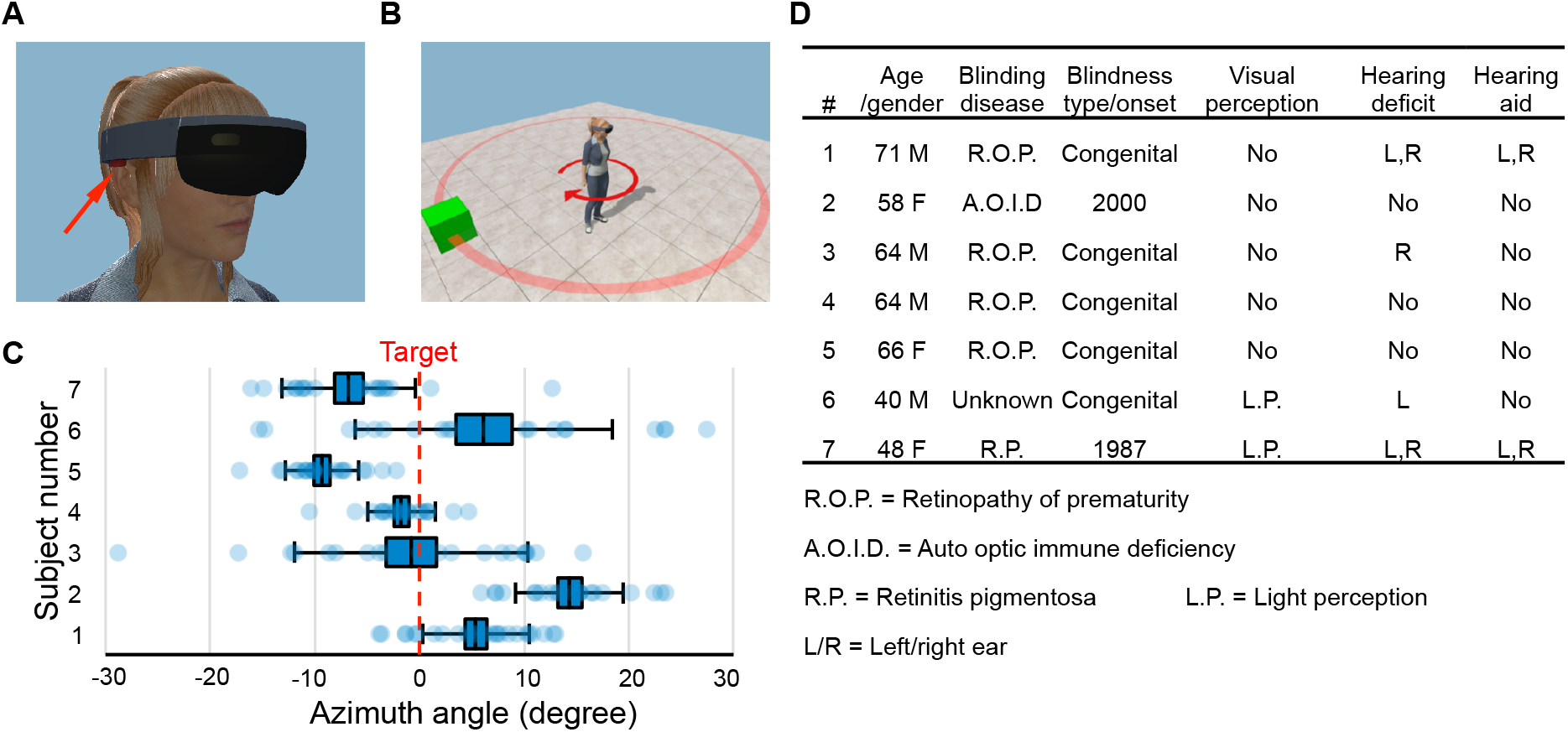
Hardware platform and object localization task. (**A**) The Microsoft HoloLens wearable augmented reality device. Arrow points to one of its stereo speakers. (**B**) In each trial of the object localization task, the target (green box) is randomly placed on a circle (red). The subject localizes and turns to aim at the target. (**C**) Object localization relative to the true azimuth angle (dashed line). Box denotes s.e.m., whiskers s.d. (**D**) Characteristics of the 7 blind subjects. See also Figures 1–4 – Source Data.

Our design principle is to give sounds to all relevant objects in the environment. Unlike most efforts at scene sonification (Bujacz and Strumillo, 2016; Csapo and Wersenyi, 2013), our system communicates through natural language. Each object in the scene can talk to the user with a voice that comes from the object’s location. The voice’s pitch increases as the object gets closer. The user actively selects which objects speak through several modes of control (Fig. S2): In Scan mode, the objects call out their names in sequence from left to right, offering a quick overview of the scene. In Spotlight mode, the object directly in front speaks, and the user can explore the scene by moving the head. In Target mode, the user selects one object that calls repeatedly at the press of a clicker. In addition obstacles and walls emit a hissing sound as the user gets too close (Fig. S2).

### Human subject tests

After a preliminary exploration of these methods we settled on a fixed experimental protocol and recruited seven blind subjects (Fig. 1D). Subjects heard a short explanation of what to expect, then donned the HoloLens and launched into a series of four fully automated tasks without experimenter involvement. No training sessions were provided, and all the data were gathered within a 2-hour visit.

### Object localization

Here we tested the user’s ability to localize an augmented reality sound source (Fig. 1B). A virtual object placed randomly at a 2 m distance from the subject called out “box” whenever the subject pressed a clicker. The subject was asked to orient the head towards the object and then confirm the final choice of direction with a voice command. All subjects found this a reasonable request and oriented surprisingly well, with an accuracy of 3–12 degrees (standard deviation across trials, Fig. 1C). Several subjects had a systematic pointing bias to one or the other side of the target (−9 to +13 deg, Fig. 1C), presumably related to hearing deficits (Fig. 1D), but no attempt was made to correct for this bias. These results show that users can accurately localize the virtual voices generated by HoloLens, even though the software used a generic head-related transfer function without customization.

### Spatial memory

Do object voices help in forming a mental image of the scene (Lacey, 2013) that can be recalled for subsequent decisions? A panel of 5 virtual objects was placed in the horizontal plane 2 m from the subject, spaced 30 degrees apart in azimuth (Fig. 2A). The subject scanned this scene actively using the Spotlight mode for 60 s. Then the object voices were turned off and we asked the subject to orient towards the remembered location of each object, queried in random order. All subjects performed remarkably well, correctly recalling the arrangement of all objects (Figs. 2B, S3B) with just one error (1/28 trials). Even the overall scale of the scene and the absolute positions of the objects were reproduced well from memory, to an average accuracy of ∼15 deg (rms deviation from true position, Figs. 2C-D). In a second round we shuffled the object positions and repeated the task. Here 3 of the subjects made a mistake, presumably owing to interference with the memory formed on the previous round. Sighted subjects who inspected the scene visually performed similarly on the recall task (Fig. S3B). These experiments suggest that active exploration of object voices builds an effective mental representation of the scene that supports subsequent recall and orientation in the environment. Whether the prosthesis also produces a subjective feeling that resembles “seeing” remains to be determined; this may emerge only after long-term use.

**Figure 2.**
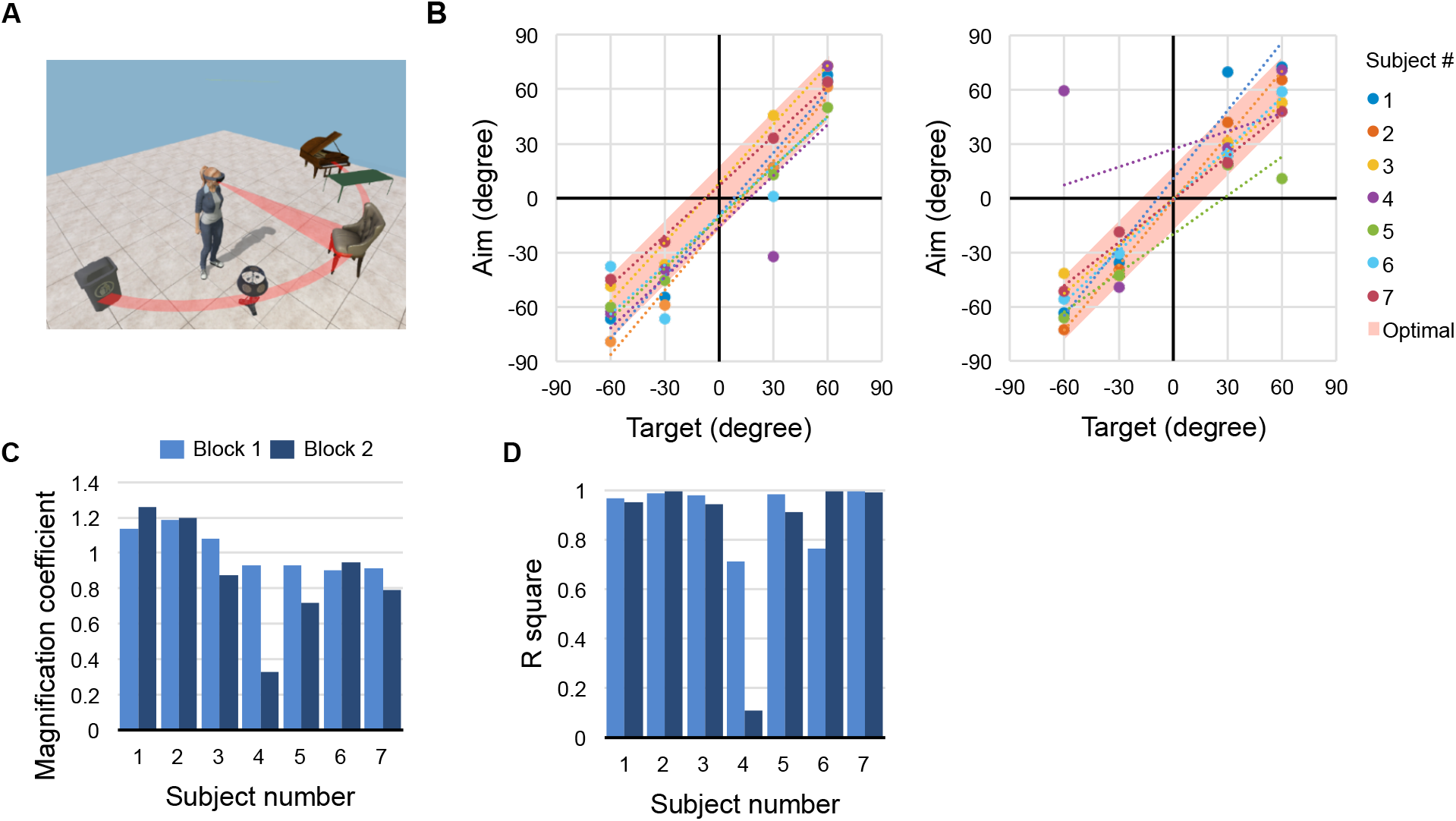
Spatial Memory Task. (**A**) Five objects are arranged on a half-circle; the subject explores the scene, then reports the recalled object identities and locations. (**B**) Recall performance during blocks 1 (left) and 2 (right). Recalled target angle potted against true angle. Shaded bar along the diagonal shows the 30 deg width of each object; data points within the bar indicate perfect recall. Dotted lines are linear regressions. (**C**) Slope and (**D**) correlation coefficient for the regressions in panel (**B**). See also Figures 1–4 – Source Data.

### Direct navigation

Here the subject was instructed to walk to a virtual chair, located 2 m away at a random location (Fig. 3A). In Target mode the chair called out its name on every clicker press. All subjects found the chair after walking essentially straight-line trajectories (Figs. 3B-C, S4). Most users followed a two-phase strategy: first localize the voice by turning in place, then walk swiftly towards it (Figs. S4D-E). On rare occasions (∼5 of 139 trials) a subject started walking in the opposite direction, then reversed course (Fig. S4C), presumably owing to ambiguities in azimuthal sound cues (McAnally and Martin, 2014). Subject 7 aimed consistently to the left of the target (just as in the task of Fig. 1) and thus approached the chair in a spiral trajectory (Fig. 3C). Regardless, for all subjects the average trajectory was only 11–25% longer than the straight-line distance (Figs. 3E, S4A).

**Figure 3.**
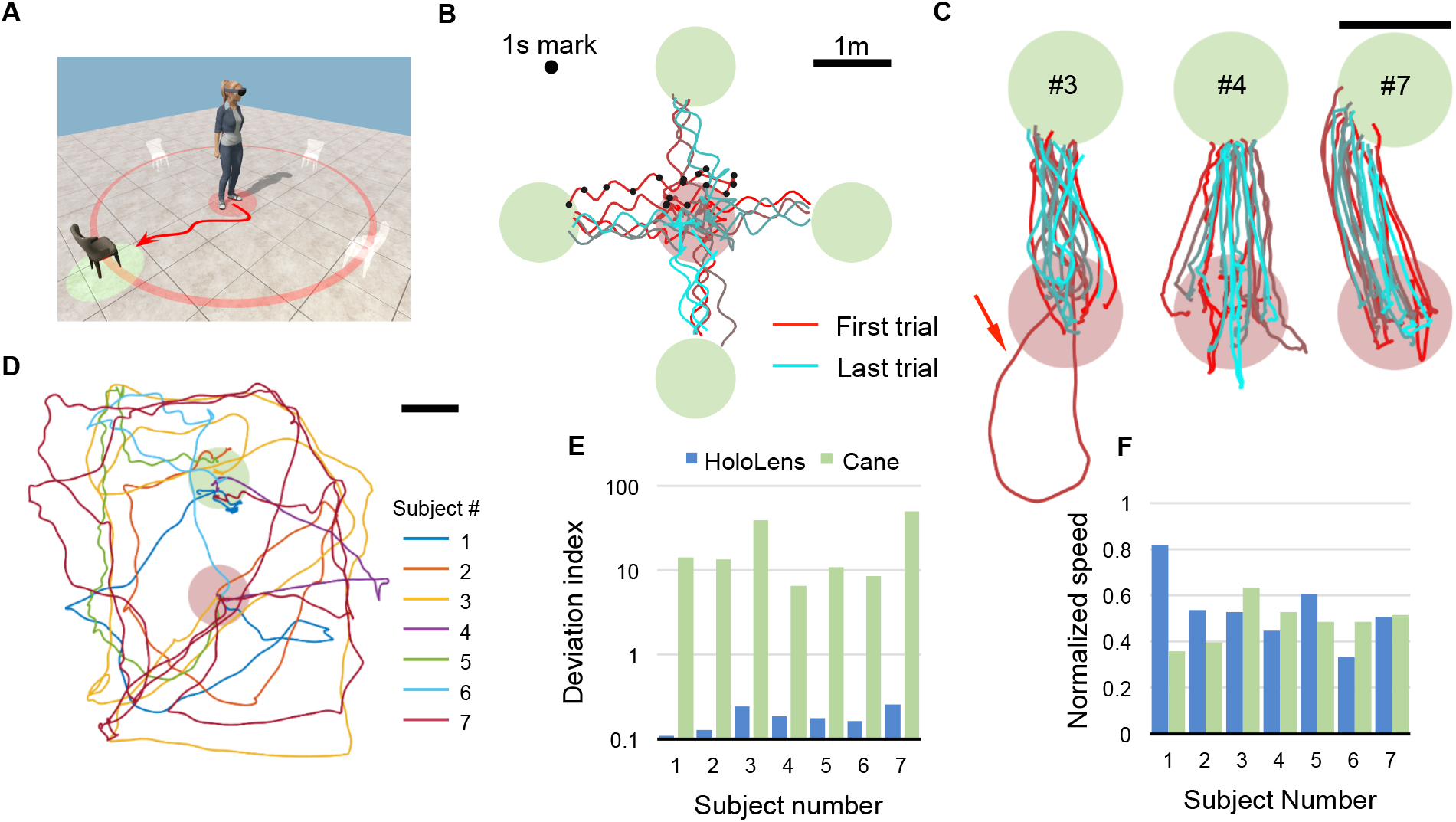
Direct Navigation Task. (**A**) For each trial a target chair is randomly placed at one of four locations. The subject begins in the starting zone (red shaded circle), follows the voice of the chair, and navigates to the target zone (green shaded circle). (**B**) All raw trajectories from one subject (#6) including 1-s time markers. Oscillations from head movement are filtered out in subsequent analysis. (**C**) Filtered and aligned trajectories from all trials of 3 subjects (#3, 4, 7). Arrow highlights a trial where the subject started in the wrong direction. (**D**) Trajectories of subjects performing the task with only a cane and no HoloLens. (**E**) Deviation index, namely the excess length of the walking trajectory relative to the shortest distance between start and target. Note logarithmic axis and dramatic difference between HoloLens and Cane conditions. (**F**) Speed of each subject normalized to the free-walking speed. See also Figures 1–4 – Source Data and Figure 3 – Source Data File Trajectories.

**Figure 4.**
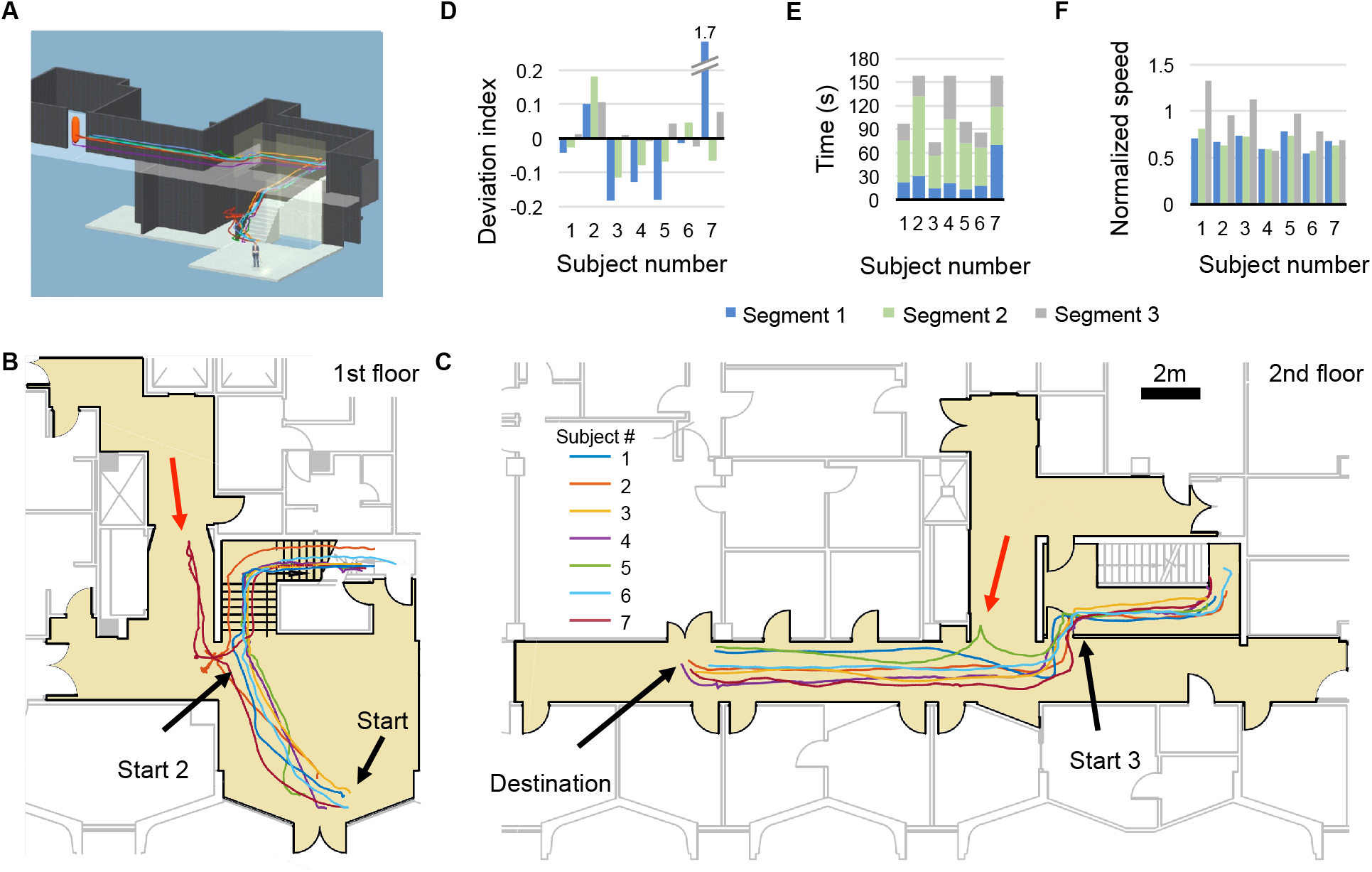
Long range guided navigation task. (**A**) 3D reconstruction of the experimental space with trajectories from all subjects overlaid. (**B** and **C**) 2D floor plans with all first trial trajectories overlaid. Trajectories are divided into 3 segments: lobby (Start – Start 2), stairwell (Start 2 – Start 3), and hallway (Start 3 – Destination). Red arrows indicate significant deviations from the planned path. (**D**) Deviation index (as in Fig. 3E) for all segments by subject. Outlier corresponds to initial error by subject 7. Negative values indicate that the subject cut corners relative to the virtual guide. (**E**) Duration and (**F**) normalized speed of all the segments by subject. See also Figures 1–4 – Source Data and Figure 4 – Source Data File Trajectories.

For comparison, we asked subjects to find a real chair in the same space using only their usual walking aid (Fig. 3D). These searches took on average 8 times longer and covered 13 times the distance needed with the prosthesis. In a related series of experiments we encumbered the path to the target with several virtual obstacles. Using the alarm sounds our subjects weaved through the obstacles without collision (Fig. S5D). Informal reports from the subjects confirmed that steering towards a voice is a natural function that can be performed automatically, leaving attentional bandwidth to process other real and augmented sounds from the environment.

### Long range guided navigation

If the target object begins to move as the subject follows its voice, it becomes a “virtual guide”. We designed a guide that follows a precomputed path and repeatedly calls out “follow me”. The guide monitors the subject’s progress, and stays at most 1 m ahead of the subject. If the subject strays off the path the guide stops and waits for the subject to catch up. The guide also offers warnings about impending turns or a flight of stairs. To test this design, we asked subjects to navigate a campus building that had been pre-scanned by the HoloLens (Figs. 4A, S6). The path led from the ground-floor entrance across a lobby, up two flights of stairs, around several corners and along a straight corridor, then into a 2nd floor office (Figs. 4B-C). The subjects had no prior experience with this part of the building. They were told to follow the voice of the virtual guide, but given no assistance or coaching during the task.

All seven subjects completed the trajectory on the first attempt (Figs. 4B-C, Supplementary Movie S1). Subject 7 transiently walked off course (Fig. 4B), due to her left-ward bias (Figs. 1C, 3C), then regained contact with the virtual guide. On a second attempt this subject completed the task without straying. On average, this task required 119 s (range 73–159 s), a tolerable investment for finding an office in an unfamiliar building (Fig. 4E). The median distance walked by the subjects was 36 m (Fig. 4D), slightly shorter (∼1%) than the path programmed for the virtual guide, because the subjects can cut corners (Fig. 4C). The subjects’ speed varied with difficulty along the route, but even on the stairs they proceeded at ∼60% of their free-walking speed (Fig. 4F). On arriving at the office, one subject remarked “That was fun! When can I get one?” (see Supplementary Observations).

### Technical extensions

A complete sensory prosthesis must acquire knowledge about the environment and then communicate that knowledge to the user. So far we have focused primarily on the second task, the interface to the user. For the acquisition of real-time knowledge, computer vision will be an important channel. Tracking and identifying objects and people in a dynamic scene still presents a challenge (see Supplementary Materials), but the capabilities for automated scene analysis are improving at a remarkable rate, propelled by interests in autonomous vehicles (Jafri et al., 2014; Verschae and Ruiz-del-Solar, 2015). We have already implemented real-time object naming for items that are easily identified by the HoloLens, such as standardized signs and bar codes (Sudol et al., 2010) (Figs. S5A-B). Furthermore, we have combined these object labels with a scan of the environment to compute in real time a navigable path around obstacles towards any desired target (Fig. S5C, Supplementary Movies S2–S3).

## Discussion

Some components of what we implemented can be found in prior work (Botezatu et al., 2017; Ribeiro et al., 2012; Wang et al., 2017). Generally assistive devices have been designed to perform one well-circumscribed function, such as obstacle avoidance or route finding (Loomis et al., 2012; Roentgen et al., 2008). Our main contribution here is to show that augmented reality with object voices offers a natural and effortless human interface on which one can build many functionalities that collectively come to resemble seeing.

Our developments so far have focused on indoor applications to allow scene understanding and navigation. Blind people report that outdoor navigation is supported by many services (access vans, GPS, mobile phones with navigation apps) but these all fall away when one enters a building (Karimi, 2015). The present cognitive prosthesis can already function in this underserved domain, for example as a guide in a large public building, hotel, or mall. The virtual guide can be programmed to offer navigation options according to the known building geometry. Thanks to the intuitive interface, naïve visitors could pick up a device at the building entrance and begin using it in minutes. In this context, recall that our subjects were chosen without prescreening, including cases of early and late blindness and various hearing deficits (Fig. 1D): They represent a small but realistic sample of the expected blind user population.

The functionality of this prosthesis can be enhanced far beyond replacing vision, by including information that is not visible. As a full service computer with online access, the HoloLens can be programmed to annotate the scene and offer ready access to other forms of knowledge. Down the line one can envision a device that is attractive to both blind and sighted users, with somewhat different feature sets, which may help integrate the blind further into the community. By this point we expect that the reader already has proposals in mind for enhancing the cognitive prosthesis. A hardware/software platform is now available to rapidly implement those ideas and test them with human subjects. We hope that this will inspire developments to enhance perception for both blind and sighted people, using augmented auditory reality to communicate things that we cannot see.

“Seeing is knowing what is where by looking” (Marr, 1982). The prosthesis described here conveys “what” by the names of objects and “where” by the location from where each object calls. “Looking” occurs when the user actively requests these calls. The principal reason sighted people rely on vision much more than audition is that almost all objects in the world emit useful light signals almost all the time, whereas useful sound signals from our surroundings are few and sporadic. Our prosthesis can change this calculus fundamentally, such that all the relevant objects emit useful sounds. It remains to be seen whether prolonged use of such a device will fundamentally alter our perception of hearing to where it feels more like seeing.

## Materials and Methods

### General implementation

The hardware platform for the cognitive prosthesis is the Microsoft HoloLens Development Edition, without any modifications. This is a self-contained wearable augmented reality (AR) device that can map and store the 3D mesh of an indoor space, localize itself in real time, and provide spatialized audio and visual display (Hoffman, 2016). We built custom software in Unity 2017.1.0f3 (64-bit) with HoloToolkit-Unity-v1.5.5.0. The scripts are written in C# with MonoDevelop provided by Unity. The experiments are programmed on a desktop computer running Windows 10 Education and then deployed to Microsoft HoloLens. The software is versatile enough to be easily deployed to other hardware platforms, such as AR enabled smart phones.

### User interface

Before an experiment, the relevant building areas are scanned by the experimenter wearing the HoloLens, so the system has a 3D model of the space ahead of time. For each object in the scene the system creates a voice that appears to emanate form the object’s location, with a pitch that increases inversely with object distance. Natural spatialized sound is computed based on a generic head-related transfer function (Wenzel et al., 1993); nothing about the software was customized to individual users. Object names and guide commands are translated into English using the text-to-speech engine from HoloToolkit. The user provides input by moving the head to point at objects, pressing a wireless Clicker, using hand gesture commands or English voice commands.

In addition to instructions shown in the main body of the article, non-spatialized instructions are available at the user’s request by voice commands. The user can use two voice commands (e.g. “direction”, “distance”) to get the direction of the current object of interest or its distance. Depending on the mode, the target object can be the object label of user’s choice (Target Mode) or the virtual guide. “Turn-by-turn” instructions can be activated by voice commands (e.g. “instruction”). The instruction generally consists of two parts, the distance the user has to travel until reaching the current target waypoint, and the turn needed to orient to the next waypoint.

### Experimental design

All results in the main report were gathered using a frozen experimental protocol, finalized before recruitment of the subjects. The tasks were fully automated, with dynamic instructions from the HoloLens, so that no experimenter involvement was needed during the task. Furthermore we report performance of all subjects on all trials gathered this way. Some incidental observations and anecdotes from subject interviews are provided in Supplementary Observations. All procedures involving human subjects were reviewed and approved by the Institutional Review Board at Caltech. All subjects gave their informed consent to the experiments, and where applicable to publication of videos that accompany this article.

### Measurement

Timestamps are generated by the internal clock of the HoloLens. The 6 parameters of the subject’s head location and orientation are recorded at 5 Hz from the onset to the completion of each trial in each task. All performance measures are derived from these time series. Localization errors of the HoloLens amount to <4 cm (Liu et al., 2018), which is insignificant compared to the distance measures reported in our study, and smaller than the line width in the graphs of trajectories in Figures 3 and 4.

### Task design

*Task 1, object localization:* In each trial, a single target is placed 1 m from the subject at a random azimuth angle drawn from a uniform distribution between 0 and 360 degrees. To localize the target, the subject presses the Clicker to hear a spatialized call from the target. After aiming the face at the object the subject confirms via a voice command (“Target confirmed”). When the location is successfully registered, the device plays a feedback message confirming the voice command and providing the aiming error. The subject was given 10–15 practice trials to learn the interaction with the prosthesis, followed by 21 experimental trials. To estimate the upper limit on performance in this task, two sighted subjects performed the task with eyes open: this produced a standard deviation across trials of 0.31 and 0.36 degrees, and a bias of 0.02 and 0.06 degrees. That includes instrumentation errors as well as uncertainties in the subject’s head movement. Note that these error sources are insignificant compared to the accuracy and bias reported in Figs 1 and 2.

*Task 2, spatial memory:* This task consists of an exploration phase in which the subject explores the scene, followed by a recall phase with queries about the scene. Five objects are placed two meters from the subject at azimuth angles of –60°, –30°, 0°, 30°, 60° from the subject’s initial orientation. Throughout the experiment, a range between –7.5° and 7.5° in azimuth angle is marked by “sonar beeps” to provide the subject a reference orientation. During the 60 s exploration phase, the subject uses “Spotlight Mode”: This projects a virtual spotlight cone of 30° aperture around the direction the subject is facing and activates object voices inside this spotlight. Typically subjects scan the virtual scene repeatedly, while listening to the voices. In the recall phase, “Spotlight Mode” is turned off and the subject performs 4 recall trials. For each recall trial the subject presses the Clicker, then a voice instruction specifies which object to turn to, the subject faces in the recalled direction, and confirms with a voice command (“Target confirmed”). The entire task was repeated in two blocks that differed in the arrangement of the objects. The object sequence from left to right was “piano”, “table”, “chair”, “lamp”, “trash bin” (block 1), and “trash bin”, “piano”, “table”, “chair”, “lamp” (block 2). The center object is never selected as a recall target because 0° is marked by sonar beeps and thus can be aimed at trivially.

*Task 3, direct navigation:* In each trial, a single chair is placed at 2 m from the center of the arena at an azimuth angle randomly drawn from four possible choices: 0°, 90°, 180°, 270°. To start a trial, the subject must be in a starting zone of 1 m diameter in the center. During navigation, the subject can repeatedly press the Clicker to receive a spatialized call from the target. The trial completes when the subject arrives within 0.5 m of the center of the target. Then the system guides the subject back to the starting zone using spatialized calls emanating from the center of the arena, and the next trial begins. Subjects performed 19–21 trials. All blind subjects moved freely without cane or guide dog during this task.

To measure performance on a comparable search without the prosthesis, each subject performed a single trial with audio feedback turned off. A real chair is placed at one of the locations previously used for virtual chairs. The subject wears the HoloLens for tracking and uses a cane or other walking aid as desired. The trial completes when the subject touches the target chair with a hand. All blind subjects used a cane during this silent trial.

*Task 4, long range guided navigation:* The experimenter defined a guide path of ∼36 m length from the first-floor lobby to the second-floor office by placing 9 waypoints in the pre-scanned environment. In each trial the subject begins in a starting zone within 1.2 m of the first waypoint, and presses the Clicker to start. A virtual guide then follows the trajectory and guides the subject from the start to the destination. The guide calls out “follow me” with spatialized sound every 2 s, and it only proceeds along the path when the subject is less than 1 m away. Just before waypoints 2–8, a voice instruction is played to inform the subject about the direction of turn as well as approaching stairs. The trial completes when the subject arrives within 1.2 meters of the target. Voice feedback (“You have arrived”) is played to inform the subject about arrival. In this task all blind subjects used a cane.

*Free walking:* To measure the free walking speed, we asked subjects to walk for 20 m in a straight line in an unobstructed hallway using their preferred walking aid. Subjects 1 and 2 used a guide dog, the others a cane.

### Data analysis and visualization

MatLab 2017b (Mathworks) and Excel (Microsoft) were used for data analysis and visualization. Unity 5.6.1f1 was used to generate 3D cartoons of experiments and to visualize 3D trajectories. Photoshop CC 2017 was used for overlaying trajectories on floor plans.

*Aiming:* In task 1 and 2, aiming errors are defined as the difference between the target azimuth angle and the subject’s front-facing azimuth angle. In task 2, to correct for the delay of voice command registration, errors are measured at 1 s before the end of each trial.

*Trajectory smoothing:* The HoloLens tracks its wearer’s head movement, which includes lateral movements perpendicular to the direction of walking. To estimate the center of mass trajectory of the subject we applied a moving average with 2 s sliding window to the original trajectory.

*Length of trajectory and deviation index:* In the directed navigation task and the long range guided navigation task, we computed the excess distance traveled by the subject relative to an optimal trajectory or the guide path. The deviation index, *DI*, is defined as

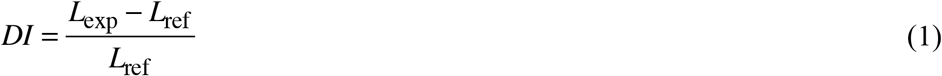

where *L*_exp_ is the length of the trajectory measured by experiment and *L*_ref_ is the length of the reference trajectory. A value near 0 indicates that the subject followed the reference trajectory well.

In the direct navigation task, we divided each trial into an orientation phase where the subject turns the body to face the target, and a navigation phase where the subject approaches the target. We calculated head orientation and 2D distance to target in each frame, and marked the onset of the navigation phase when the subject’s distance to target changed by 0.3 m. Note that with this criterion the navigation phase includes the occasional trajectory where the subject starts to walk in the wrong direction. In this task *L*_ref_ is defined as the length of the straight line from the subject’s position at the onset of the navigation phase to the nearest point of the target trigger zone.

In the long range guided navigation task, *L*_ref_ is the length of the guide trajectory. Due to variability in placing waypoints and tracking, the length of guide trajectories varied slightly across subjects (*L*_ref_ = 36.4 ± 0.7 m, mean ± sd). Negative *DI* values are possible in this task if the subject cuts corners of the guide trajectory.

*Speed:* Speed is calculated frame-by-frame using the displacements in the filtered trajectories. For the long range guided navigation task, which includes vertical movements through space, the speed of translation is computed in 3 dimensions, whereas for the other tasks that occur on a horizontal plane we did not include the vertical dimension. For all tasks, we estimated walking speed by the 90th percentile of the speed distribution, which robustly rejects the phases where the subject chooses an orientation. The normalized speed is obtained by dividing this value by the free walking speed.

## Acknowledgments

Supported by a grant from the Shurl & Kay Curci Foundation.

## Data and materials availabilit

Data and code that produced the figures are available in the Source Data Files.

## Supplementary Materials

Supplementary Methods

Supplementary Observations

Fig. S1. Process of scene sonification.

Fig. S2. Obstacle avoidance utility and active scene exploration modes.

Fig. S3. Object localization task and mental imagery task supplementary data (related to Figs. 1 and 2).

Fig. S4. Direct navigation task extended data (related to Fig. 3).

Fig. S5. Additional experimental functions.

Fig. S6. Guided navigation trajectories (related to Fig. 4).

Movie S1. Long range navigation (Fig. 4), Subject 6.

Movie S2. Automatic wayfinding explained (Fig. S5).

Movie S3. Automatic wayfinding (Fig. S5), Point of View during navigation.

Source Data Files for Figs 1–4.

## Supplementary Materials

### This file includes

Supplementary Methods

Supplementary Observations

Supplementary Figures S1 to S6

Captions and external links for Movies S1 to S3

Links to Source Data Files for Figures 1 to 4.

### Supplementary Methods

#### Voice Control

In addition to the Clicker, subjects can also use natural language (e.g. English) as input to the system. Two subsystems of voice input are implemented: 1) keyword recognition (PhraseRecognitionSystem) monitors in the background what the user says, detects phrases that match the registered keywords, and activates corresponding functions on detection of keyword matches. 2) dictation (DictationRecognizer) records what the user says and converts it into text. The first component enables subjects to confirm their aiming in the object localization task and mental imagery task with the voice command “target confirmed”. It also enables the experimenter to control the experiment at runtime.

Keywords and their functions are defined through adding keywords to the keyword manager script provided by HoloToolkit and editing their responses. The KeywordRecognizer component starts at the beginning of each instance of the application and runs in the background throughout the instance of the application except for the time period in which dictation is in use.

To allow users to create object labels, the DictationRecognizer provided by HoloToolkit is used to convert natural language spoken by the user to English text. Due to the mutual exclusivity, KeywordRecognizer is shut down before DictationRecognizer is activated, and restarted after the dictation is finished.

#### Soundscape Editing

*Adding Object Labels:* Users can manually add object labels at runtime with voice commands (e.g. “record label”). Each call of the object label adding function instantiates an object label where the user is aiming. The system listens to what the user says for a certain period of time (e.g. 3 s), and converts the speech into text at the same time.

When the dictation finishes, the converted text will be read, the user is asked for a confirmation of the recorded text, and a timer starts. The user uses voice commands (e.g. “confirm”) to confirm the addition of the object label and the content of the label before the timer reaches a certain time limit. At the same time, the object label list is updated to include the new object label. If no confirmation is received and time runs out, the newly created object label is deleted.

In addition to manual labeling, a computer vision based toolkit Vuforia SDK (v6.1.17 distributed by PTC Inc. of Boston, Massachusetts) is used for recognizing and tracking objects using the forward-facing camera on the HoloLens. We trained it to recognize a restroom sign and to create a virtual object (label) on top of it (Fig. S5A). The created virtual object persists even when the HoloLens can no longer see the original sign (Fig. S5B).

*Deleting Object Labels:* To delete an object label, the user first chooses the object label to be deleted as the object of interest in the Target Mode, and then uses voice commands (e.g. “delete label”) to delete the chosen object label. Immediately after the deletion of an object label, the list of object labels is updated.

*Moving Object Labels:* In Developer Mode, objects can be relocated by the user. An object label is in the placing mode when the user aims at it and clicks on it. When it enters the placing mode, the object label floats at a fixed distance in front of the user. The user clicks again to re-anchor the object label in the environment.

#### Automated Wayfinding

In addition to hand-crafting paths, we implemented automated wayfinding by taking advantage of Unity’s runtime NavMesh “baking” which calculates navigable areas given a 3D model of the space. At runtime, we import and update the 3D mesh of the scanned physical space and use it to bake the 3D mesh. When the user requests guided navigation, a path from the user’s current location to the destination of choice is calculated. If the calculated path is valid, the virtual guide guides the user to the destination using the computer-generated path.

#### Cost of the system

The hardware platform used in the research – Microsoft HoloLens Development Edition – currently costs $3000. Several comparable AR goggles are in development, and one expects their price to drop in the near future. In addition, smart phones are increasingly designed with AR capabilities, although they do not yet match the HoloLens in the ability to scan the surrounding space and localize within it.

#### Battery and weight

The current HoloLens weighs 579 g. Like all electronic devices, this will be further miniaturized in the future. The current battery supports our system functions for 2–5 h, sufficient for the indoor excursions we envision in public buildings, led by the “virtual guide”. A portable battery pack can extend use to longer uninterrupted sessions.

#### Tracking robustness

While in most indoor scenarios that we have tested the tracking of HoloLens was reliable and precise, we have encountered occasional loss of tracking or localization errors. This occurs particularly when the environment lacks visual features such as a narrow space with white walls.

#### Dynamic scenes

To maintain a smooth user experience, the HoloLens updates its internal model of the real-world space every few seconds. This computational bottleneck limits its capability of mapping highly dynamic scenes, such as a busy store with many customers walking around. However, with increasing computational power packed into mobile devices and the development of more efficient scene understanding algorithms this performance is expected to improve accordingly. There is a large software industry dedicated to solving these problems of real time scene understanding, and the cognitive prosthesis will be able to exploit those developments.

#### Extensions

Because this cognitive prosthesis is largely defined by software its functionalities are very flexible. For example, the diverse recommendations from subjects noted above (Supplementary Observations) can be implemented in short order. In addition one can envision hardware extensions by adding peripherals to the computer. For example a haptic belt or vest could be used to convey collision alarms (Adebiyi et al., 2017), thus leaving the auditory channel open for the highly informative messages.

### Supplementary Observations

Here we report incidental observations not planned in the frozen protocol, and comments gathered from blind subjects in the course of the experiments.

*Subject 1:* During navigation with the virtual guide says “seems to me the ‘follow me’ sound means keep going straight”. Thinks addition of GPS services could make the system useful outdoors as well. Suggests experimenting with bone conduction headphones. Offers us 1 hour on his radio show.

*Subject 2:* During direct navigation says “pitch change [with distance] was informative”. During navigation with the virtual guide says “‘Follow me’ was too much information”. Prefers to follow the explicit turn instructions. She could then transmit those instructions to her guide dog.

*Subject 3:* In addition to object voices, he likes instructions of the type ‘keep going forward for xx meters’. During a previous visit using a similar system he commented on possible adoption by the blind community: “I could see people spending in 4 figures for [something] light and reliable, and use it all the time”. Also supports the concept of borrowing a device when visiting a public building or mall. Devices in the form of glasses would be better, preferably light and thin. “Use the computing power of my phone, then I don’t have to carry anything else.” Likes the external speakers because they don’t interfere with outside sound. Finds it easy to localize the virtual sound sources.

*Subject 4:* After navigation with the virtual guide says “That was fun. When can I get one?” Primarily used the ‘follow me’ voice, and the cane to correct for small errors. Reports that the turn instructions could be timed earlier (this is evident also in movie S1). On a previous visit using a similar system: “I’m very excited about all of this, and I would definitely like to be kept in the loop”. Also suggests the system could be used in gaming for the blind.

*Subject 5:* During navigation with the virtual guide realized she made a wrong turn (see Fig. 4C) but the voice made her aware and allowed her to correct. Reports that the timing of turn instructions is a little off.

*Subject 6:* After all tasks says “That was pretty cool” and “The technology is there.”

*Subject 7:* On the second trial with the virtual guide reports that she paid more attention to the ‘follow me’ sound (she strayed temporarily on the first trial, Fig. 4B). Wonders whether the object voices will be strong enough in a loud environment.

**Supplementary Figure S1.**
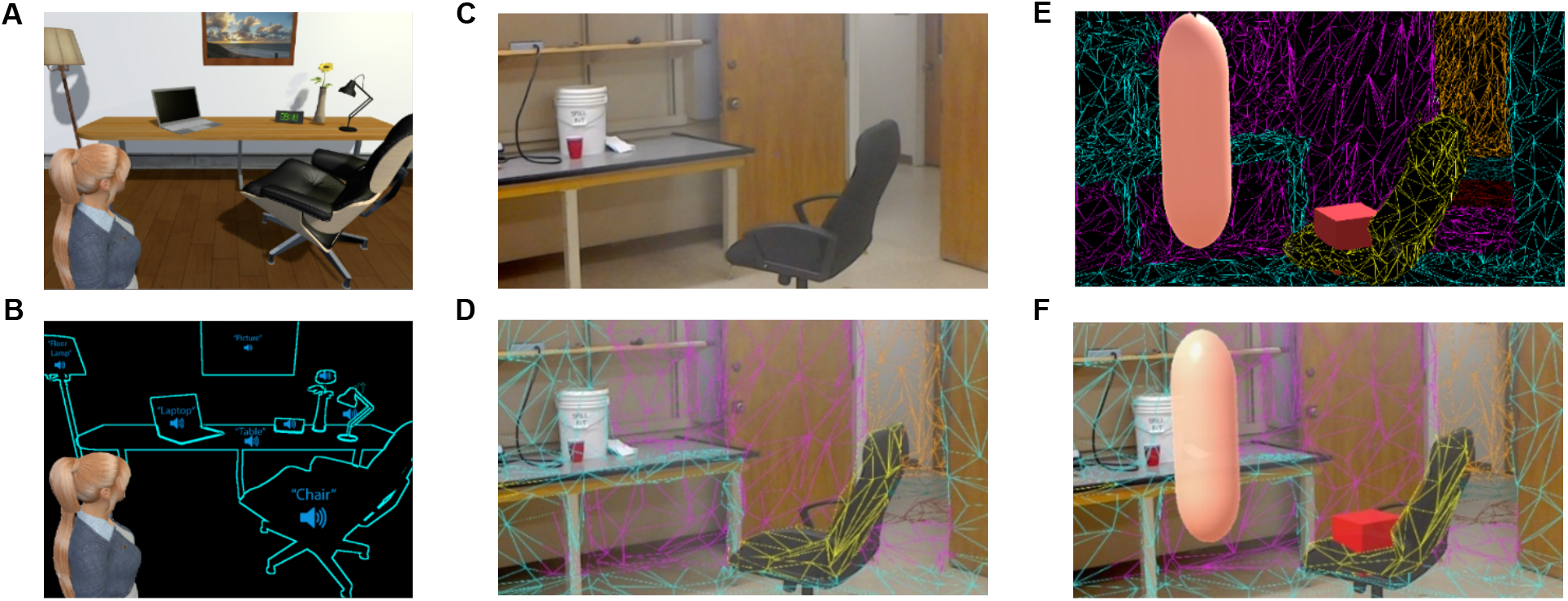
Process of scene sonification. The acquisition system should parse the scene (**A**) into objects and assign each object a name and a voice (**B**). In our study this was accomplished by a combination of the HoloLens and the experimenter. The HoloLens scans the physical space (**C**) and generates a 3D mesh of all surfaces (**D**). In this digitized space (**E**) the experimenter can perform manipulations such as placing and labeling virtual objects, computing paths for navigation, and animating virtual guides (**F**). Because of the correspondence established in D, these virtual labels are tied to the physical objects in real space.

**Supplementary Figure S2.**
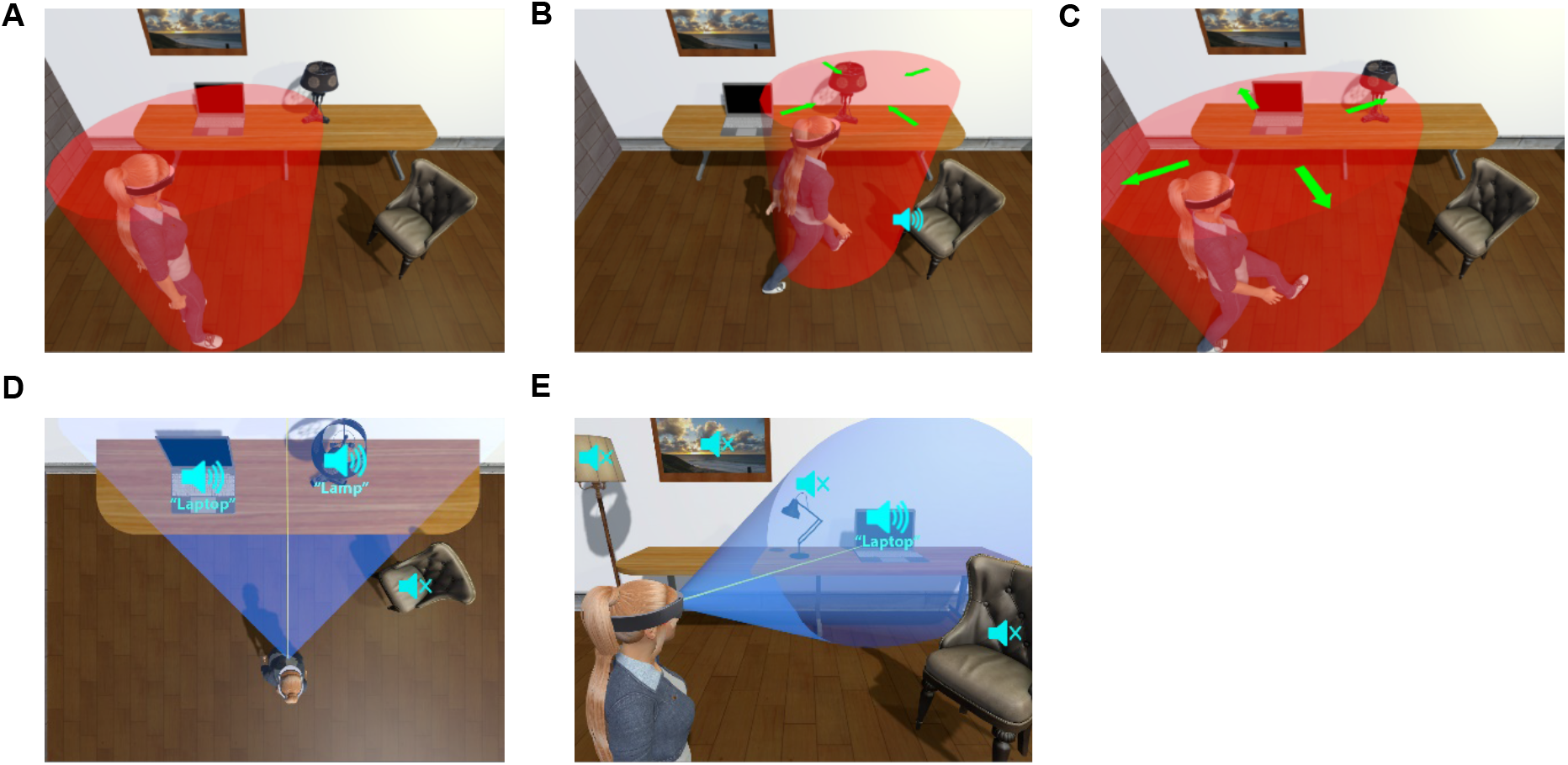
Obstacle avoidance utility and active scene exploration modes. (**A** to **C**) An object avoidance system is active in the background at all times. Whenever a real scanned surface or a virtual object enters a danger volume around the user (red in **A**), a spatialized warning sound is emitted from the point of contact (**B**). The danger volume expands automatically as the user moves (**C**), so as to deliver warnings in time. (**D** to **E**) Active exploration modes. In Scan mode (**D**) objects whose azimuthal angles fall in a certain range (e.g. between –60 and +60 deg) call themselves out from left to right. In Spotlight mode (**E**) only objects within a narrow cone are activated, and the object closest to the forward-facing vector calls out.

**Supplementary Figure S3.**
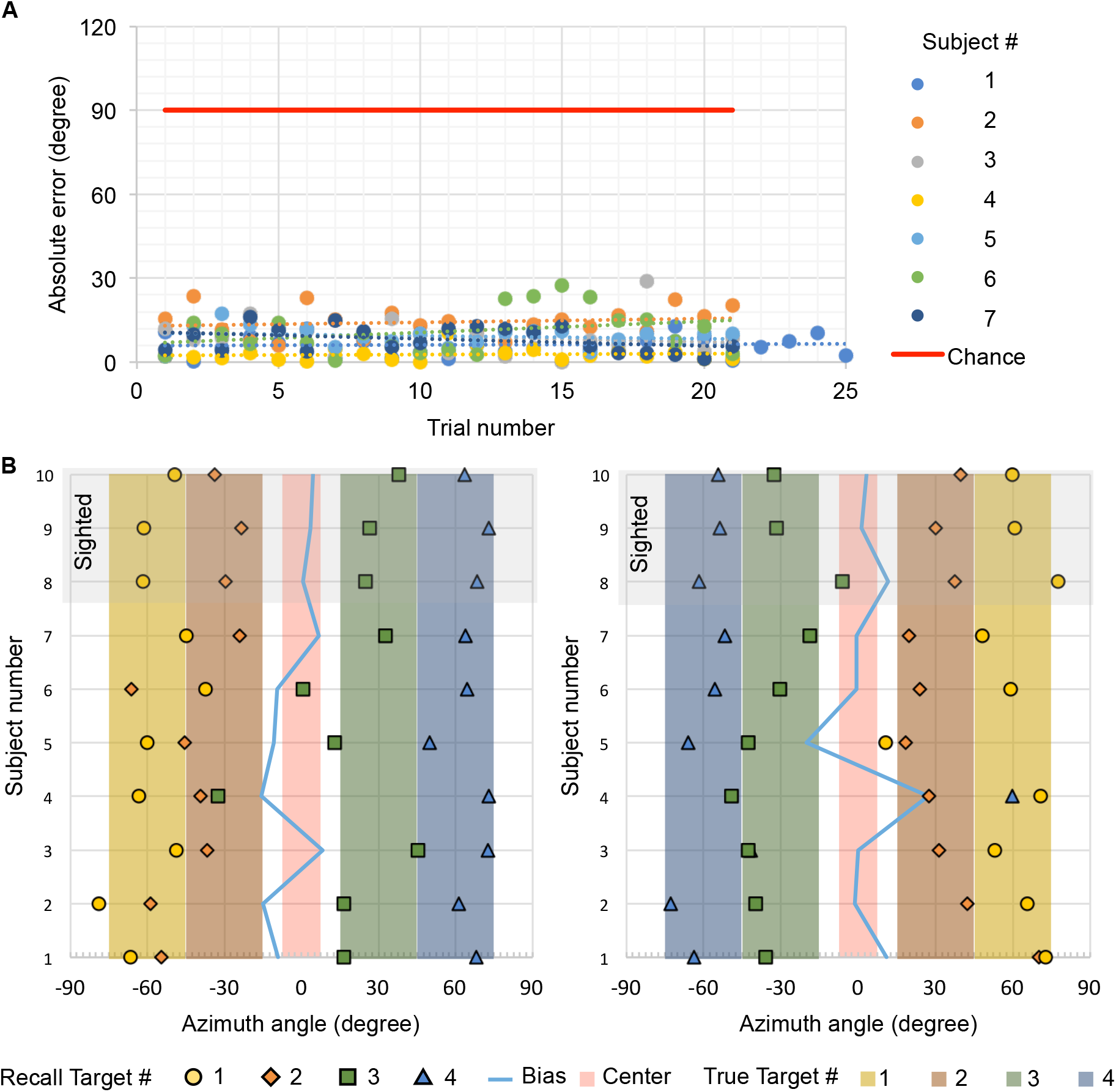
Object localization task and mental imagery task supplementary data (related to Figs. 1 and 2). (**A**) Absolute error of object localization (Fig. 1) by trials. Chance level is 90 deg. (**B**) Spatial memory data (Fig. 2) from block 1 (left) and 2 (right) by subject. Shaded areas indicate the true azimuthal extent of each object. Markers indicate recalled location. Most recalled locations overlap with the true extent of the object. Subjects 8–10 were normally sighted and performed the exploration phase using vision.

**Supplementary Figure S4.**
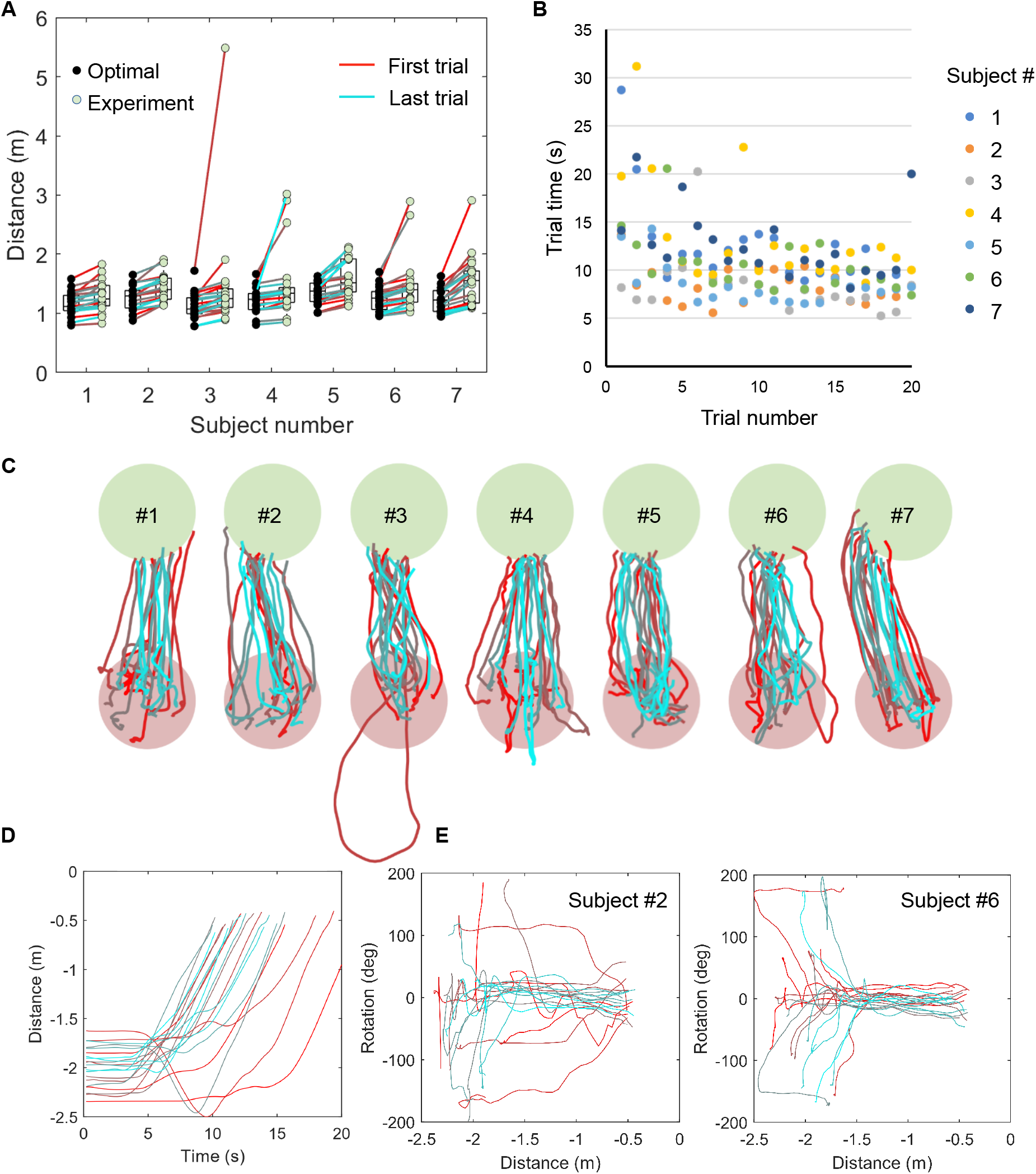
Direct navigation task extended data (related to Fig. 3). Trial distance (**A**) and trial duration (**B**) for the first 20 trials of all subjects A modest effect of practice on task duration can be observed across all subjects (**B**). (**C**) Low-pass filtered, aligned trajectories of all subjects. In most trials, subjects reach the target with little deviation. (**D**) Dynamics of navigation, showing the distance to target as a function of trial time for one subject. (**E**) Head orientation vs distance to target for two subjects. Note subject 6 begins by orienting without walking, then walks to the target. Subject 2 orients and walks at the same time, especially during early trials.

**Supplementary Figure S5.**
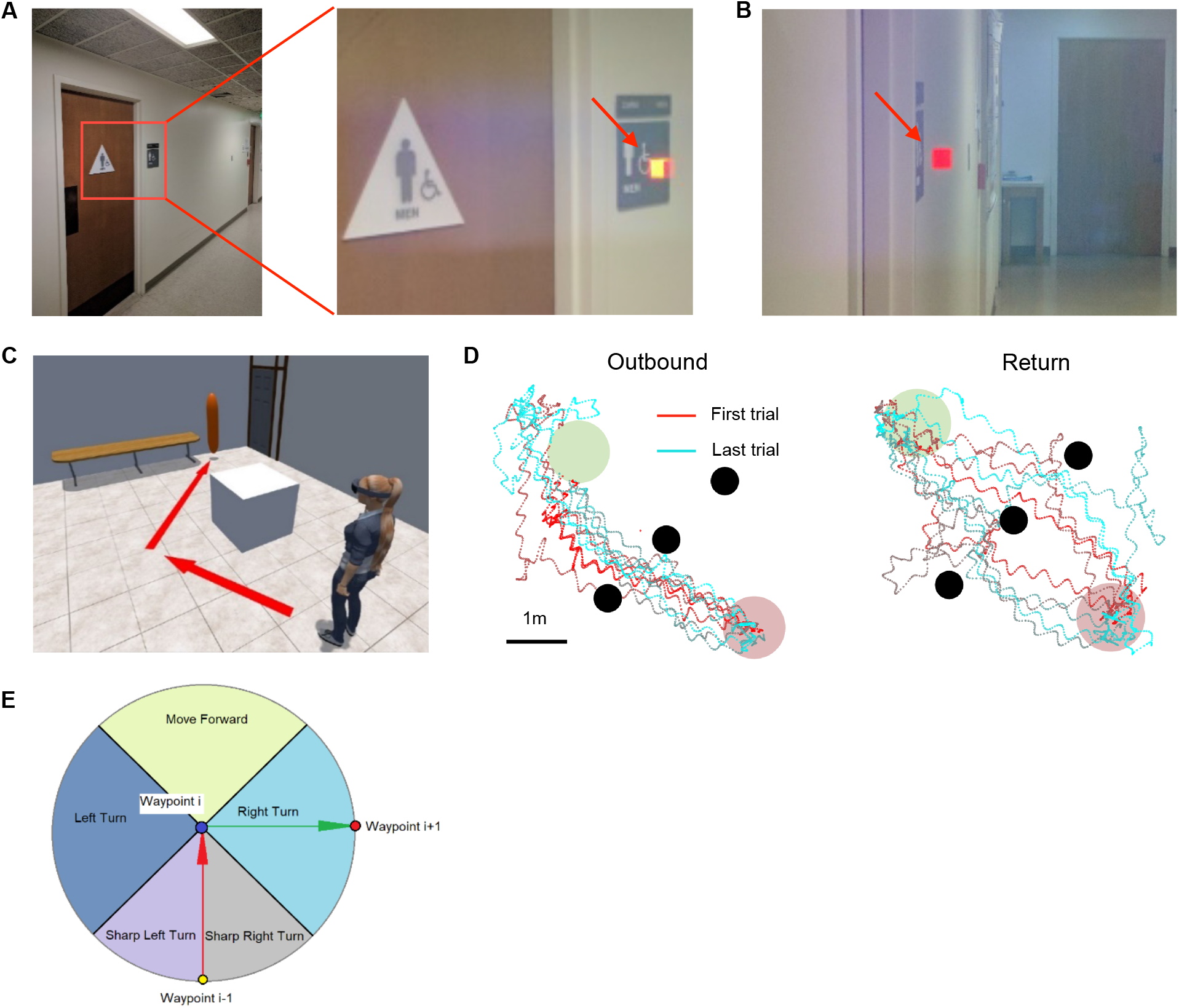
Additional experimental functions. (**A** to **B**) Automated sign recognition using computer vision. Using Vuforia software (https://www.vuforia.com/) the HoloLens recognizes a men’s room sign (**A**, image viewed through HoloLens) and installs a virtual object (cube, arrow) next to the sign. (**B**) This object persists in the space even when the sign is no longer visible. (**C**) Automated wayfinding. The HoloLens generates a path to the target (door) that avoids the obstacle (white box). Then a virtual guide (orange balloon) can lead the user along the path. See Movies S2—S3. (**D**) Navigation in the presence of obstacles. The subject navigates from the starting zone (red circle) to an object in the target zone (green circle) using calls emitted by the object. Three vertical columns block the path (black circles), and the subject must weave between them using the obstacle warning system. Raw trajectories (no filtering) of a blind subject (#5) are shown during outbound (left) and return trips (right), illustrating effective avoidance of the columns. This experiment was performed with an earlier version of the apparatus built around the HTC Vive headset. (**E**) Orienting functions of the virtual guide. In addition to spatialized voice calls the virtual guide may also offer turning commands towards the next waypoint. In the illustrated example, the instruction is “in x meters, turn right.”

**Supplementary Figure S6.**
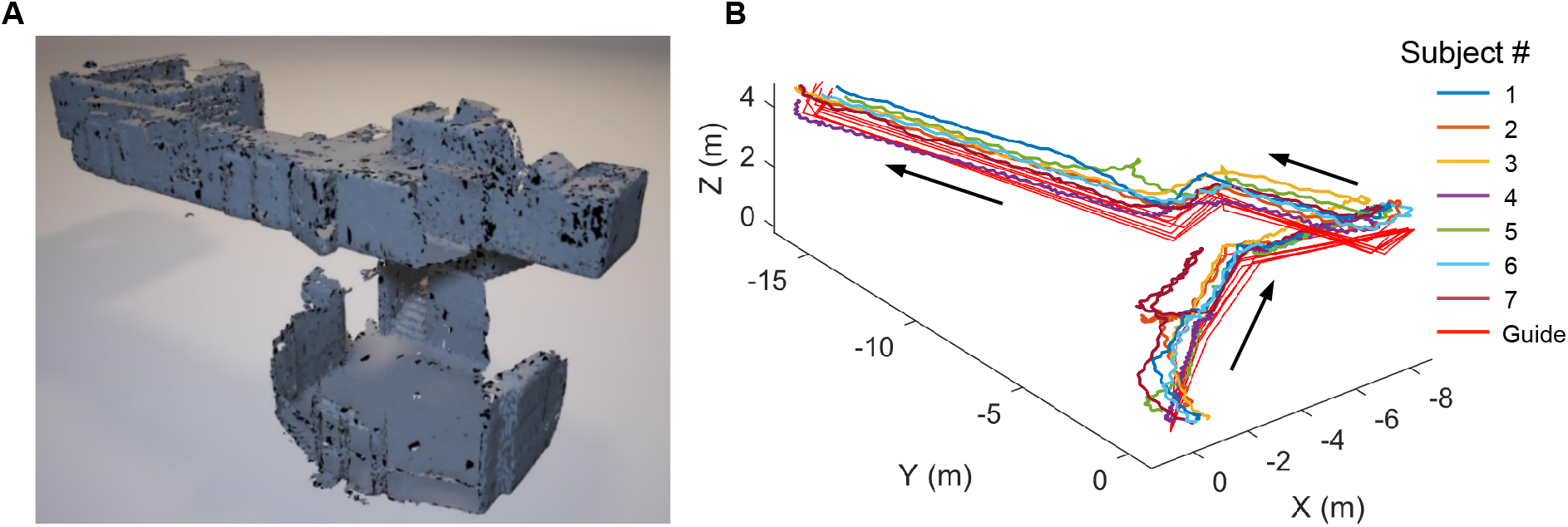
Guided navigation trajectories (related to Fig. 4). (**A**) 3D model of the experimental space as scanned by the HoloLens. (**B**) Subject and guide trajectories from the long range guided navigation task. Note small differences between guide trajectories across experimental days, owing to variations in detailed waypoint placement.

**Movie S1. Long range navigation (Fig. 4), Subject 6.**

https://drive.google.com/open?id=1v5Wdbi2WWXAMQXyLmVU6ogWDQfpyINQu

**Movie S2. Automatic wayfinding explained (Fig. S5).**

https://drive.google.com/open?id=1B6kx89Ce35w_aNTc-Q4ExGhA3RRLfrXm

**Movie S3. Automatic wayfinding (Fig. S5), Point of View during navigation.**

https://drive.google.com/open?id=1R72kbfHsbqxuxEcbLj0KfISyRAzXWKH_

**Figures 1–4 – Source Data File:**

https://drive.google.com/open?id=1Sy_Ky2d0GIkyoPiH23xvbrGIpJETvdIh

**Figure 3 – Source Data File Trajectories:**

https://drive.google.com/open?id=1gCb0hqMA0Uol9QcLFlhyz3F5hrNFhEvs

**Figure 4– Source Data File Trajectories:**

https://drive.google.com/open?id=18nJxqtqZ3irNVMVR5CvQkx4JcyE7GUrV

